# Short-term exposure to high-water temperature causes a shift in the microbiome of the common aquarium sponge *Lendenfeldia chondrodes*

**DOI:** 10.1101/2020.01.10.901363

**Authors:** Sergio Vargas, Laura Leiva, Gert Wörheide

## Abstract

Marine sponges harbor diverse microbiomes that contribute to their energetic and metabolic needs. Although numerous studies on sponge microbial diversity exist, relatively few focused on sponge microbial community changes under different sources of environmental stress. In this study, we assess the impact of elevated seawater temperature on the microbiome of cultured *Lendenfeldia chondrodes*, a coral reef sponge commonly found in marine aquaria. *Lendenfeldia chondrodes* exhibits high thermal tolerance showing no evidence of tissue damage or bleaching at 5 °C above control water temperature (26 °C). High-throughput sequencing of the bacterial 16S rRNA V4 region revealed a response of the microbiome of *L. chondrodes* to short-term exposure to elevated seawater temperature. Shifts in abundance and richness of the dominant bacterial phyla found in the microbiome of this species, namely Proteobacteria, Cyanobacteria, Planctomycetes, and Bacteroidetes, characterized this response. The observed resilience of *L. chondrodes* and the responsiveness of its microbiome to short-term increases in seawater temperature suggest that this holobiont may be capable of acclimating to anthropogenic-driven sublethal environmental stress via a re-accommodation of its associated bacterial community. This sheds a new light on the potential for resilience of some sponges to increasing surface seawater temperatures and associated projected regime shifts in coral reefs.

## Introduction

By the end of this century, moderate climate change scenarios predict the surface temperature of the globe to be over 1.5 °C higher than in 1850-1900, and the oceans are expected to keep warming throughout the 21st century [1]. Extreme events, such as the marine heatwave recorded on the Great Barrier Reef in 2016, are expected to become more common under those scenarios [1], and their detrimental impact on coastal ecosystems will undoubtedly continue to cause significant shifts in the species composition of marine communities [e.g. 2].

An increase in seawater temperature of one to two degrees over maximum summer temperatures approaches the upper tolerance level of important coastal ecosystem engineers, like corals [3] and some sponges tested to date [4–9]. The reef sponge *Rhopaloeides odorabile*, for instance, showed signs of necrosis after 24 hours of exposure to seawater temperatures of 33 °C, only about two to four degrees above the mean highest summer temperature at the site of collection [4]. At this temperature, the bacterial community associated with *R. odorabile* suffers irreversible compositional shifts likely caused by a “breakdown” of the sponge symbiotic functions and the colonization of the sponge by opportunistic, potentially pathogenic bacteria [4]. Simister et al. [10] reported similar results. These authors found a shift in the bacterial community associated with *R. odorabile* after exposing this sponge to a seawater temperature of 32 °C. In both experiments, the authors reported a highly stable microbiome at temperatures below that upper limit [4, 10].

Beside *R. odorabile*, several reports of a highly stable sponge microbiome at temperatures below a specific threshold exists. The microbiome of the Australian reef sponge *Ianthella basta* is stable across a latitudinal gradient of over 1000 km and at temperatures up to 31 °C, with microbiome changes only evident after exposure of the sponges to lethal water temperatures [>32 °C, 5]. Also, the microbiome of the Mediterranean species *Ircinia fasciculata* and *I. oros* appears to be resistant to sublethal heat stress [11]. However, once the water temperature exceeds an upper limit, the microbiome of *I. fasciculata* changes, usually preceding the onset of disease and, eventually, the death of the affected sponges [9]. Perhaps unexpectedly, the microbiome of the boreal sponge *Geodia barreti* was also not affected by exposure to acute thermal stress (i.e., 5 °C above “normal” environmental seawater temperature) for up to 14 days [12]. Thus, the resilience of the sponge microbiome to sublethal thermal stress does not appear to be determined by the geographic area where the sponges occur.

Throughout its ∼600 million years long evolutionary history [13, 14] Phylum Porifera has evolved to become a significant part of benthic communities worldwide [15], occurring from the tropics to the polar regions in habitats ranging from the eulittoral to the hadal zone [15–17] and playing essential roles in nutrient cycling in these ecosystems [15, 18–20]. The diverse functional roles and ecological flexibility of sponges partly result from their diverse microbiomes [21–23], and although many sponge species have highly stable microbiomes regardless of the environment [24, 25], environmental changes can trigger compositional shifts in sponge microbiomes [26] that can affect the fitness of these holobionts. Thus, investigating the effect of environmental change on the sponge microbiome is pivotal to our understanding of the consequences of climate change upon sponge holobionts and the benthic communities where these organisms occur.

*Lendenfeldia chondrodes*, a common (blue) aquarium sponge, is an easy-to-culture, promising (cyano)sponge holobiont model [27]. Here, we characterized the microbiome of *L. chondrodes* using high-throughput 16S rRNA sequencing and examined the effect of a rapid, short-term increase in sea surface temperature (SST) on the diversity and community composition of the microbiome of this species.

## Material and Methods

### Heat shock experiments

We cultured purple and green morphs of *Lendenfeldia chondrodes* (Suppl. Figure 1) in the aquaria of the Molecular and Geobiology & Paleobiology Laboratory of the Dept. of Earth- and Environmental Science of the Ludwig-Maximilians Universität München. For the heat shock experiments, we cut circular sections of the sponges with a medical biopsy puncher (0.5 cm diameter) and kept them under control conditions for one week before transferring them to the experimental tanks. We assigned at random six experimental tanks (20cm x 20cm, 6L) to either the control or the heat-stress treatments, and assigned at random two *L. chondrodes* explants per color morph to each tank. In total, we allocated six replicates of each color morph (i.e., purple or green) to each treatment. We gradually increased the seawater temperature of the treatment tanks over five days, from ∼25.4 °C to 30 °C (i.e., approximately one Celsius degree per day), using a TetraTec HT 25 heating device. We then exposed the sponges to 30 °C SST for two days before terminating the experiment. The water temperature of the control tanks was ∼25.4 °C over the whole experimental period. At the end of the experiment, we fixed all explants in 99% ethanol and kept the samples at −20 °C until further processing.

**Figure 1.**
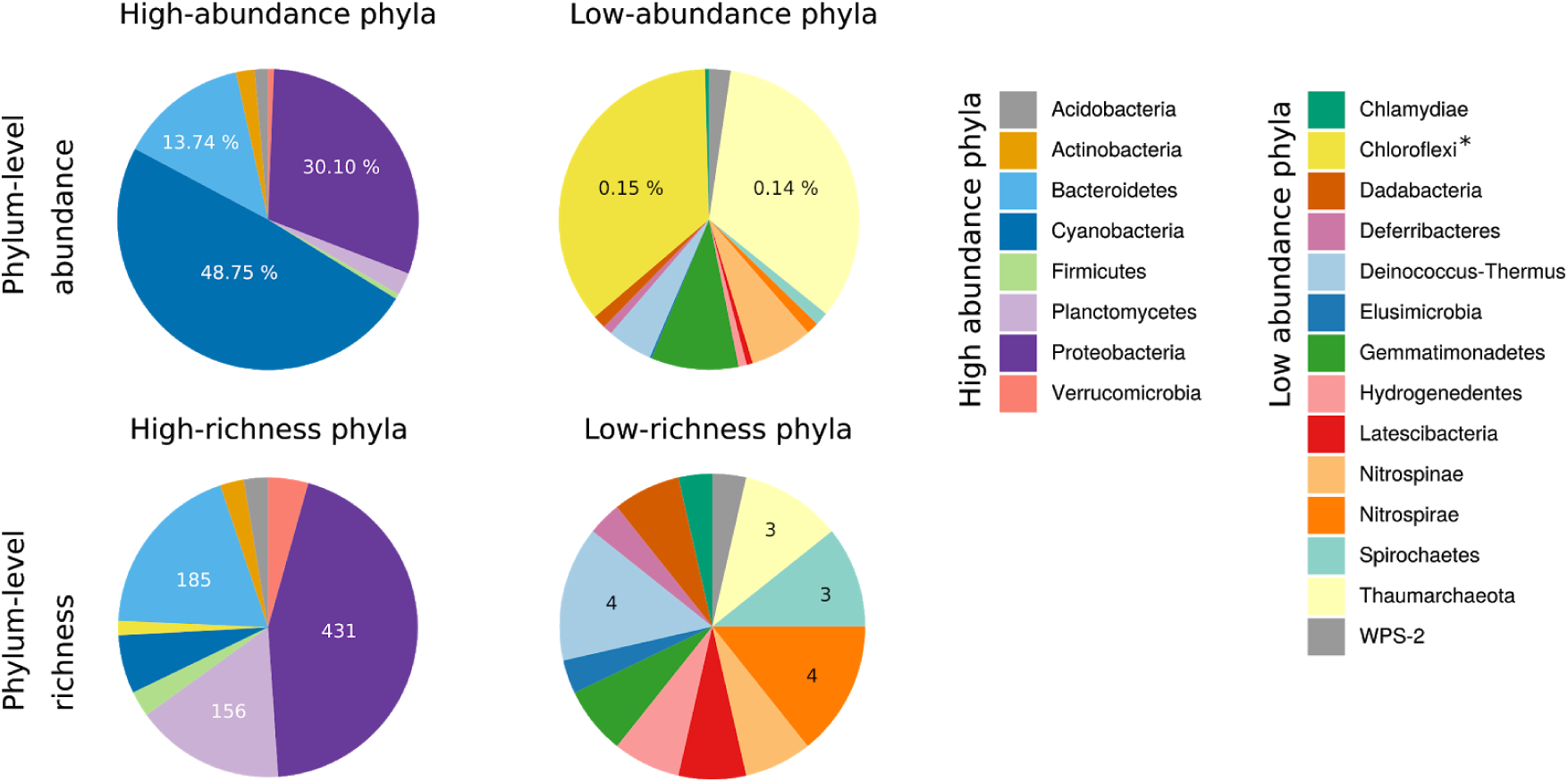
Abundance and richness of bacterial phyla associated with the common aquarium sponge *Lendenfeldia chondrodes*. High-abundance phyla are defined as having at least 1000 16S V4 rRNA read counts. High-richness phyla are those with at least 10 OTUs.

### High-throughput sequencing bacterial 16S rRNA V4 region of control and treated L. chondrodes

We extracted genomic DNA using the Macherey-Nagel NucleoSpin DNA extraction kit, following the protocol provided by the manufacturer, and amplified the V4 region of the 16S rRNA gene using barcoded forward (515fB) and reverse (806rB) primers as described in [28]. Before sequencing, we visualized PCR products on 1% agarose gels and extracted bands of the expected size (ca. 380bp) with the Qiagen QIAquick Gel Extraction Kit. We then quantified all gel-extracted PCR products on a Qubit 3.0 fluorometer (Life Technologies, Grand Island, NY) and diluted them, if necessary, to achieve a final concentration of 1nM before equimolar pooling. We sequenced the 16S rRNA amplicon pool on an Illumina MiniSeq in mid-output 300PE mode [see 28 for details]. We also determined the bacterial load of each extraction using real-time quantitative PCR (RT-qPCR). For this, we used the same forward (i.e., 515fB) and reverse (i.e., 806rB) primers but without the barcode+Illumina adapter extension used for the generation of Illumina-ready amplicon libraries. We use the bacterial load of each sample to provide absolute OTU counts after amplicon sequencing (see below). The reads generated are publicly available under study accession number PRJEB35927.

### Bioinformatic analysis of the 16S rRNA amplicon data

After demultiplexing, we processed the resulting sequences using vsearch [29]. Briefly, we assembled each sequence pair allowing a minimum overlap of 45bp, dereplicated within and across samples, removed singletons and chimeras, and clustered sequences at 97% similarity to derive a “raw” Operational Taxonomic Unit (OTU) counts table. We then used the RT-qPCR data on bacterial load to provide absolute counts per sample for each OTU in the OTU table. For this, briefly, we rescaled the obtained OTU counts to the total number of bacteria per uL DNA. We determined the taxonomic affiliation of all OTUs found in *L. chondrodes* using the SILVA database “Alignment, Classification and Tree Service” (accessed the 22.05.2019) and used the OTU count tables and the taxonomy annotations to determine phylum- and OTU-level richness and abundance patterns, and to fit rank-abundance dominance curves to the bacterial community associated with *L. chondrodes*. We also determined the core bacterial community of *L. chondrodes* as those bacterial OTUs present in all sequenced replicates independently of the treatment applied to them.

To assess the beta-diversity of treated *vs.* control sponges, we used non-metric multidimensional scaling (NMDS) with the Bray-Curtis distance. We also tested for significant effects of heat stress on bacterial composition using canonical correspondence analysis (DCA) with treatment (i.e., Control *vs.* Heat stress), color morph (i.e., Green *vs.* Purple) and their interaction as response variables. In order to test for a differential response to heat stress of core *vs.* non-core OTUs, we conducted separate DCAs for these bacterial groups. To better understand the dynamics of the bacterial community in response to heat stress, we identified OTUs gained or lost as a result of the treatment. For core OTUs, we also calculated log-fold changes in heated *vs.* control samples.

We used R with the packages vegan [30] and DESeq2 [31] for the analyses and deposited all scripts and raw data tables used in the project repository (https://gitlab.lrz.de/cbas/CBAS_16S).

## Results

### The microbiome of L. chondrodes is rich but dominated by a small number of bacterial associates

In total, we found 1343 OTUs affiliated to 22 bacterial phyla associated with *L. chondrodes*. Phylum Proteobacteria had the highest OTU richness (431 OTUs), followed by Bacteroidetes (185 OTUs) and Planctomycetes (156 OTUs). The remaining 19 phyla had considerable less OTUs assigned to them. For instance, we found 61 and 42 OTUs belonging to the phyla Cyanobacteria and Firmicutes, respectively, and all remaining phyla had 25 or fewer OTUs (Fig. 1). Dominance patterns differed radically from those of richness, with Cyanobacteria dominating the community and accumulating 47% of all the sequenced reads. The phyla Proteobacteria and Bacteroidetes followed in abundance with 29% and 13% of the reads, respectively. The remaining 19 phyla accounted for 11% of all the sequenced reads (Fig. 1). The core bacterial community of *L. chondrodes* is composed of 17 bacterial OTUs belonging to the phyla Acidobacteria, Actinobacteria, Bacteroidetes, Cyanobacteria, Planctomycetes, and Proteobacteria (Suppl. Table 1). These OTUs accounted for 78% of the sequenced reads across all samples.

**Table 1.** Permutation test (999 permutations) for a canonical correspondence analysis of the effect of temperature treatment, sponge color morph and their interaction on the composition of the microbiome of *Lendenfeldia chondrodes*. Terms added sequentially, first to last.

|  | Df | Chi-Square | F | Pr(>F) |
| --- | --- | --- | --- | --- |
| Treatment | 1 | 0.20026 | 1.7458 | 0.008 |
| Color morph | 1 | 0.11300 | 0.9850 | 0.492 |
| Treatment x Color morph | 1 | 0.13395 | 1.1677 | 0.191 |
| Residual | 16 | 1.83543 |  |  |

As judged by its Akaike Information Criterion (AIC), the phylum-level best fit abundance-rank dominance model was a geometric series (niche preemption model; [32] for 15 out of 20 replicates, followed by the Mandelbrot and the lognormal model in four and one replicates, respectively (Fig. 2). Fitting the Mandelbrot model to the phylum-level abundance data was challenging, and some parameters of this model could not be fit. Whenever the full model could be specified, the AIC for the Mandelbrot model was lower than that for the niche preemption model. Generally, the analysis of model deviance across replicates resulted in a non-significant difference between these models (Suppl. Table 2). At OTU-level, the best fit (i.e., lowest AIC) abundance-rank dominance models were the Zipf (15 replicates) or Mandelbrot (four replicates), with only one replicate fitting the lognormal model (Fig. 2). As in the analysis of the phylum-level abundances, an analysis of model deviance revealed that the Zipf and Mandelbrot models were not significantly different (Suppl. Table 3).

**Figure 2.**
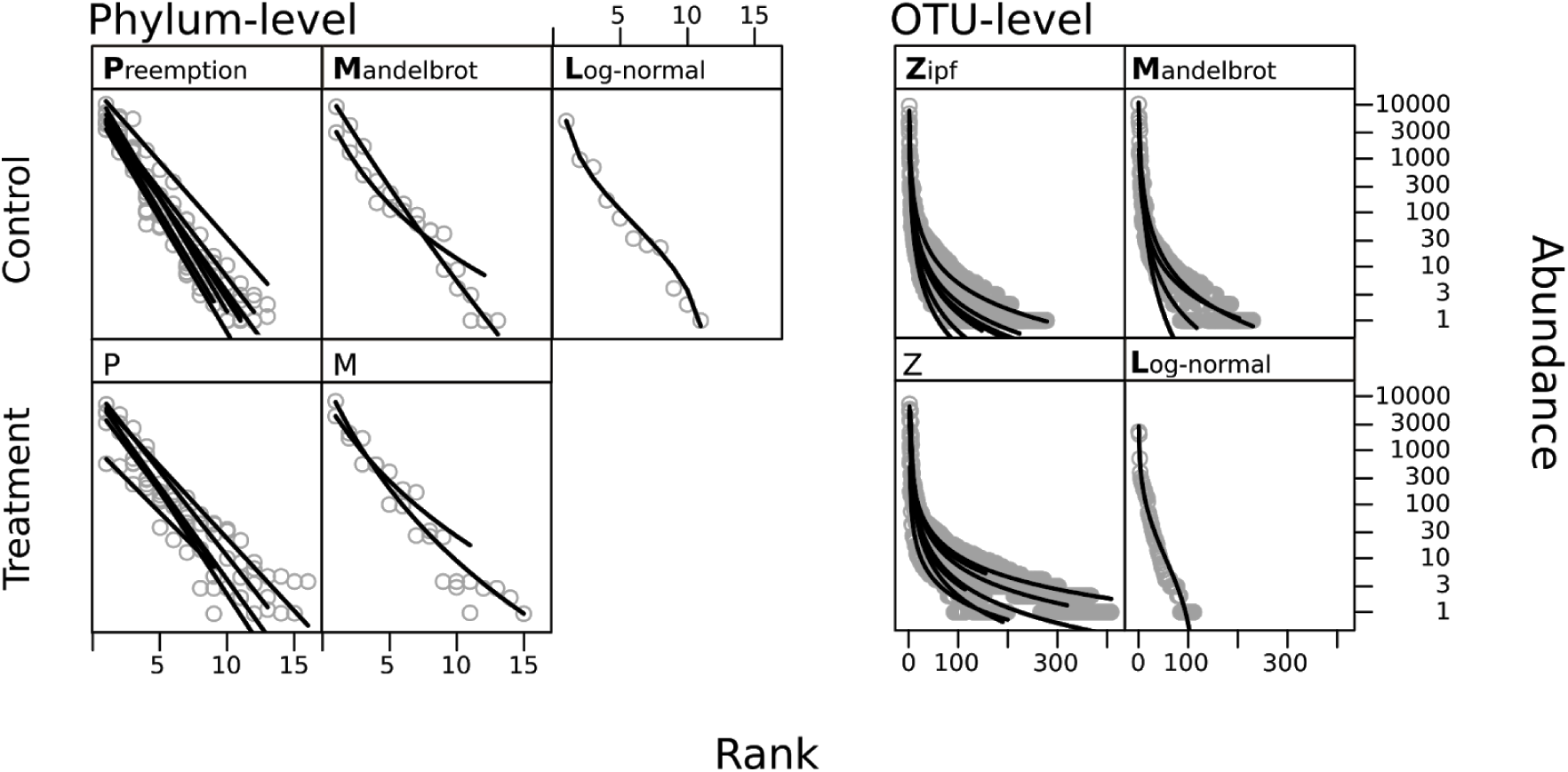
Abundance-rank dominance models fitted to Phylum- and OTU-level 16S V4 rRNA counts for control and thermal stress treatments. Lines represent sample replicates, dots are phyla/OTUs. The name of the models are above the curves, the letter in bold is used as the model abbreviation.

### Beta-diversity changes in core vs. non-core bacteria in a specific manner upon short-term exposure of L. chondrodes to heat stress

Heat stress caused a compositional change of the bacterial communities associated with *L. chondrodes*. This effect was somewhat mild in the NMDS plots (Fig. 3A) but marked in the canonical correspondence analysis, where control and treatment samples formed two distinct groups (Fig. 3B). Indeed, we found that the variable “Treatment” (i.e., heat stress vs. control temperature) had a significant effect on bacterial composition while the variable “Color morph” did not; the interaction between these two variables was not significant (Table 1). These results were independent of any filtering (e.g., including only OTUs with more than 50 reads) of the raw count OTU table we used for the analysis. However, these results differed from those we obtained using the bacterial load corrected OTU table. If we used these data, the NMDS analysis did not converge to a solution unless we filtered the OTU table to include only OTUs with more than 50 (absolute) counts. Using this (filtered) table, we also observed a significant effect of the variable “Treatment” on bacterial composition (Suppl. Table 4). In contrast to the analysis of the raw count table, we observed a significant effect of the variable “Color morph” on bacterial composition, while the interaction between these variables was still not significant (Suppl. Table 4).

**Figure 3.**
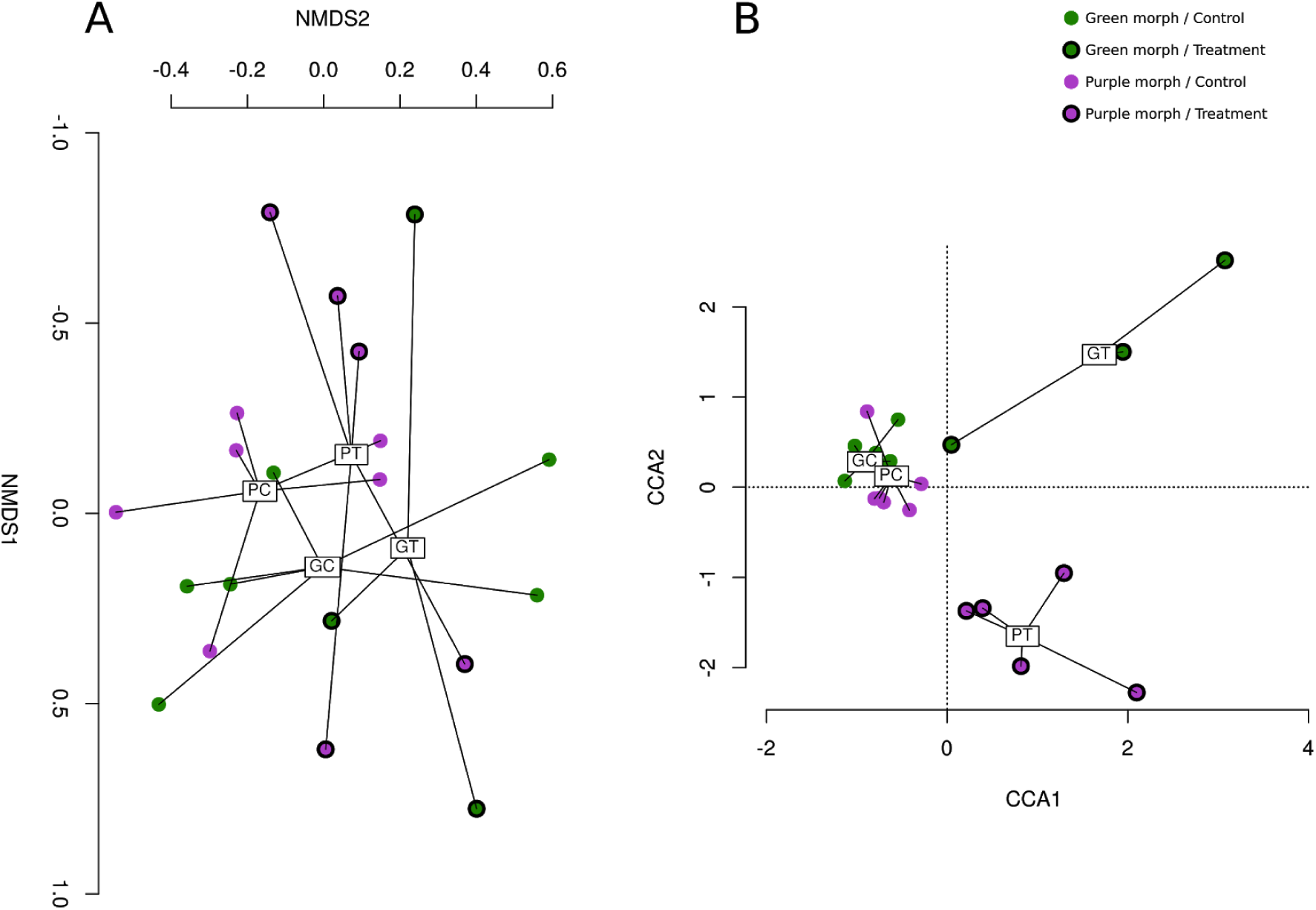
Non-metric multidimensional scaling (A) and canonical correspondence (B) analysis of the OTU-level composition of the microbiome of *Lendenfeldia chondrodes* in control vs. thermal stress treatments. GC: Green morph control samples, PC: Purple morph control samples, GT: Green morph treatment samples, PT: Purple morph treatment samples.

The analysis of core and non-core bacterial OTUs revealed a similar trend and allowed us to better dissect the effect of heat stress on both types (i.e., core and non-core) of bacterial associates (Suppl. Figure 2). Here again, we detected a significant effect of the heat stress treatment on both core and non-core bacterial associates. However, we only detected a significant effect of sponge color morph for non-core bacterial OTUs. Moreover, this group of OTUs was also affected by the interaction between thermal treatment and sponge color morph. In line with these results, we found that the heat stress treatment induced a change in the community characterized by the net loss of OTUs in treated vs. control sponges (Fig. 4).

**Figure 4.**
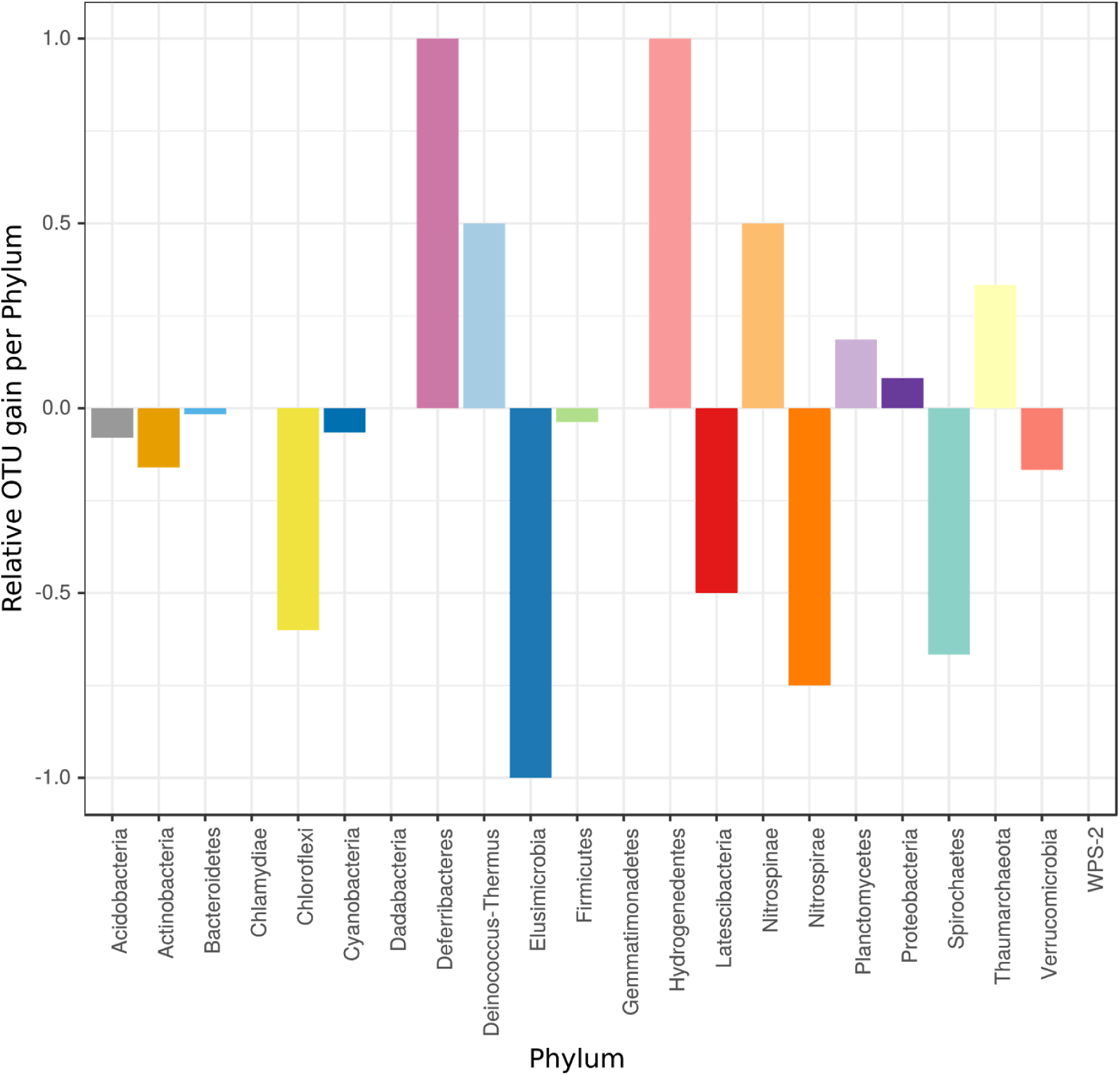
Relative OTU gain/loss per Phylum after exposure to thermal stress. Bacterial gain/loss is defined as the appearance/disappearance of an OTU in the treated samples taking as reference the composition of control samples.

Here, 11 out of 22 phyla detected in *L. chondrodes* loss OTUs, while only seven phyla gained new OTUs in response to heat stress. As expected, OTU replacement had a more notable impact on the richness of rare phyla, causing the disappearance of one bacterial phylum (Elusimicrobia) and lowering the richness of other bacterial phyla like Chloroflexi, Latescibacteria, Nitrospirae and, Spirochaetes in treated samples to less than half of the values detected in control sponges. Bacterial phyla benefiting from the heat stress treatment include Deferribacteres and Hydrogenedentes, which were only detectable in heat stress samples, and Deinococcus-Thermus and Nitrospinae, which duplicated their richness in treated vs. control sponges. Thaumarcheota also increased its richness in response to heat stress.

### Heat stress affects different members of the core bacterial community associated with L. chondrodes in a specific manner

The analysis of abundance log-fold changes of *L. chondrodes’* 17 core bacterial OTUs revealed that heat stress had a specific effect on different members of this community. This differential response was not taxon-specific as the abundance of OTUs belonging to the same phylum responded in opposite ways upon exposure to heat stress (Fig. 5). For instance, within Proteobacteria, which with ten OTUs is the richest phylum in the core community of *L. chondrodes*, three OTUs halved (i.e., log-fold change < −1) and four OTUs doubled (i.e., log-fold change > 1) their abundances. This trend was also evident in the other phyla with more than one OTU, namely Bacteroidetes and Cyanobacteria. In both phyla, one OTU increased its abundance while the abundance of the remaining OTU assigned to the phylum decreased. The observed change in OTU abundance seems to correlate with the mean overall abundance of these OTUs (Fig. 6). This pattern appears to be general for the core bacterial community. Indeed, the four most and less abundant OTUs decreased and increased their abundance in response to heat stress. Despite this, we did not find a significant correlation between abundance log fold change and mean overall OTU abundance. Exploration of the regression diagnostics revealed a single, highly influential OTU (i.e., OTU 8, Proteobacteria; Suppl. Figure 3) with a large residual (e_OTU16 = 3.24 vs. mean_e = −0.20) and Cook’s D (d_OTU16 = 0.42 vs. mean_d = 0.03). After removing this single OTU from the analysis, abundance log fold change correlated negatively (Pearson’s r = −0.54, Spearman’s rho = −0.52) with mean overall OTU abundance (Suppl. Figure 3). This correlation, which was significant independently of the correlation test used, indicates that heat stress induces rare core OTUs to increase their abundance while the abundance of the dominant core OTUs decreases.

**Figure 5.**
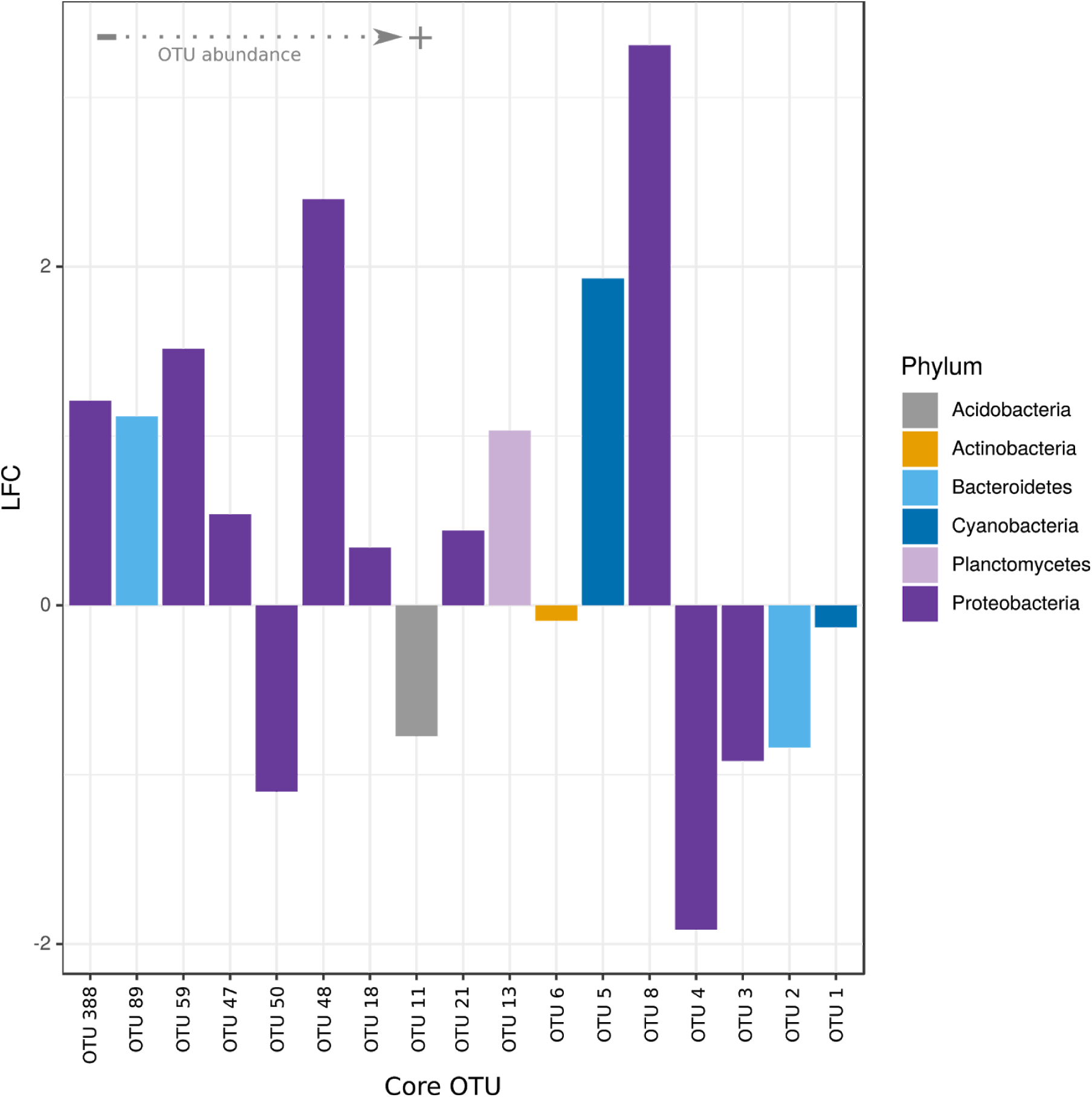
Abundance log2-fold change for core OTUs in *Lendenfeldia chondrodes* in thermal stress vs. control samples. The OTUs are ordered by the relative abundance with OTU 1 and OTU 388 being the OTUs with the highest the lowest abundance respectively.

## Discussion

Here, we provide the first characterization of the microbiome of *L. chondrodes*, a keratose sponge commonly found in salt-water aquaria around the globe [27]. With over 1300 OTUs belonging in 22 phyla, the microbial community associated with *L. chondrodes* lie well within the ranges of OTU- and Phylum-level richness (i.e., 50 to 3820 OTUs, and 13 to 34 phyla, respectively) reported in a microbiome survey of 81 sponge species [33]. The high number of OTUs assigned to Phylum Proteobacteria found in *L. chondrodes* also matches previous reports indicating that this phylum is the richest in sponge microbiomes [23]. In *L. chondrodes*, however, the phyla Bacteroidetes and Planctomycetes were also OTU-rich, which contrasts with findings in other sponges [23, 33]. These results highlight the disparity in OTU composition displayed by sponge-associated bacterial communities and further support the proposed importance of host identity in structuring their composition [33].

Sponge microbiomes harbor a mixture of host-generalists and specialists. The abundance-rank dominance models fitting the microbiome of *L. chondrodes* allow for a similar interpretation. Especially the Zipf-Mandelbrot model point to a microbiome dominated by a small set of host-generalists with few niche requirements, complemented with a low abundance but species-rich assemblage of host-specialists and transient associates with more narrow niches [34, 35]. In agreement with this, we found the “core” bacterial community associated with *L. chondrodes* to be composed of 16 OTUs that account for over 80% of the 16S rRNA reads sequenced. Reduced core communities, ranging between 7 and 20 OTUs, are common in sponge microbiomes [33].

The original interpretation of the Zipf-Mandelbrot model associates the abundance of each item with a cost, without defining precisely what the cost is [34]. In ecology, a link between the cost of a species and its trophic level exists and implies that autotrophs cost less than heterotrophs [34]. This species-cost scheme predicts a higher abundance of autotrophs in the community, in agreement with our findings in *L. chondrodes*, where a single cyanobacterial OTU accounting for almost half of the 16S rRNA reads sequenced dominated the microbiome. It seems of interest to further investigate whether the microbiomes of other cyanosponges behave similarly and fit the Zipf-Mandelbrot model, as this model allows for mechanistic interpretations of the abundance-rank data that could provide insights into the functioning of these bacterial communities.

In contrast with previous reports on several sponge species, we observed a significant compositional change in the microbiome of *L. chondrodes* upon short-term exposure to sublethal heat stress (i.e., from 27 °C to 30 °C in one week). The Caribbean sponge *Xestospongia muta* responded similarly to 12 days of exposure to sublethal heat stress [8].

In both *L. chondrodes* and *X. muta*, the observed microbiome changes did not correlate with any signs of necrosis or discoloration of the sponge tissues. The lack of evident signs of stress at sublethal water temperatures is consistent with previous experiments in other sponge species [4, 5, 10]. However, in those species, sublethal heat stress did not trigger a significant change in the composition of the microbiome. This suggests that despite its convergent nature [33], sponge microbiomes have different levels of resilience to increasing surface seawater temperatures. This indicates that anthropogenic-driven climate change and the associated projected regime shifts in coral reefs [see 36, 37] will result in loser and winner sponge species showing contrasting responses to the same environmental stimulus, and will impact the composition of sponge communities in an unpredictable manner.

The response of the *L. chondrodes* microbiome to heat stress involved a re-accommodation of both bacterial richness. OTU-rich phyla, like Proteobacteria and Planctomycetes, became even richer after the treatment, gaining the largest number of OTUs in absolute terms during the experiment. This trend was, however, not general as other OTU-rich phyla like Bacteroidetes and Cyanobacteria showed the opposite trend losing OTUs in response to heat-stress. In relative terms, however, temperature stress had a more pronounced effect among rare phyla, with some groups doubling or halving its richness and other entirely disappearing from the community. Given the lack of knowledge on this symbiotic system, and generally on the specific role of the rare microbial biosphere in the functioning of the community, linking the effects of the sudden bacterial OTU gain or loss caused by heat stress with any functional changes of the *L. chondrodes* microbiome remains challenging. The development of methods to assess changes in gene expression at the holobiont level appears thus imperative to further understand *L. chondrodes’* interactions with its microbiome.

In sponge microbiomes, density-dependent processes determine the abundance of particular OTUs in the core community [33]. Similarly, short-term exposure to sublethal heat stress affected the abundances of core OTUs in a density-dependent manner, causing an increase and a reduction of the population size of the top four most and less abundant OTUs, respectively. The variation in abundance indicates that environmental changes can either reinforce or relax the negative feedback loop controlling the population growth of the core microbiome of *L. chondrodes*. Understanding how the interaction between the intrinsic and extrinsic population factors affects the abundance of different members of the sponge microbiome is essential to determine the effect of environmental change on the fitness of specific members of the microbiome. Establishing this link will help to provide mechanistic insights into the effects of climate change on sponge holobionts and predict their response to future oceanic conditions.

## Acknowledgments

We thank Monica Pichler for her constant support during the first stages of data analysis, Peter Naumann for helping set up the aquarium experiments, and Cüneyt Çaglar and Thorsten Zimmer for their support during the execution of the heat stress experiments. LL thanks Lena Wilde and David Piatka for their constant support, feedback, and motivation. LL and SV acknowledge partial funding through the Master Program in Geobiology and Paleobiology (MGAP) of the Faculty of Geosciences and the GeoBio-Center^LMU^ of the Ludwig-Maximilians-Universität München. GW acknowledges funding through the LMU Munich’s Institutional Strategy LMUexcellent within the framework of the German Excellence Initiative. SV is indebted to N. Villalobos Trigueros, M. Vargas Villalobos, S. Vargas Villalobos, and S. Vargas Villalobos for their constant support.

## Author contributions

Conceptualization: Sergio Vargas; Data curation: Laura Leiva & Sergio Vargas; Formal analysis: Sergio Vargas; Funding acquisition: Sergio Vargas, Gert Wörheide; Investigation: Laura Leiva; Methodology: Sergio Vargas; Project administration: Sergio Vargas; Resources: Sergio Vargas, Gert Wörheide; Supervision: Sergio Vargas; Visualization: Laura Leiva & Sergio Vargas; Writing – original draft: Laura Leiva & Sergio Vargas; Writing – review & editing: Sergio Vargas, Gert Wörheide.

## References

1. IPCC (2014) Contribution of Working Groups I, II and III to the Fifth Assessment Report of the Intergovernmental Panel on Climate Change. In: Core Writing Team, R. K. Pachauri and L. A. Meyer (eds.) (ed) Climate Change 2014: Synthesis Report. IPCC, 151 pp., Geneva, Switzerland, p 151 pp

2. Hughes TP, Kerry JT, Baird AH, et al (2018) Global warming transforms coral reef assemblages. Nature 556:492–496

3. Hughes TP, Barnes ML, Bellwood DR, et al (2017) Coral reefs in the Anthropocene. Nature 546:82–90

4. Webster NS, Cobb RE, Negri AP (2008) Temperature thresholds for bacterial symbiosis with a sponge. ISME J 2:830–842

5. Luter HM, Whalan S, Webster NS (2012) Thermal and sedimentation stress are unlikely causes of brown spot syndrome in the coral reef sponge, *Ianthella basta*. PLoS One 7:e39779

6. Massaro AJ, Weisz JB, Hill MS, Webster NS (2012) Behavioral and morphological changes caused by thermal stress in the Great Barrier Reef sponge *Rhopaloeides odorabile*. J Exp Mar Bio Ecol 416-417:55–60

7. Fan L, Liu M, Simister R, et al (2013) Marine microbial symbiosis heats up: the phylogenetic and functional response of a sponge holobiont to thermal stress. ISME J 7:991–1002

8. Lesser MP, Fiore C, Slattery M, Zaneveld J (2016/2) Climate change stressors destabilize the microbiome of the Caribbean barrel sponge, *Xestospongia muta*. J Exp Mar Bio Ecol 475:11–18

9. Blanquer A, Uriz MJ, Cebrian E, Galand PE (2016) Snapshot of a Bacterial Microbiome Shift during the Early Symptoms of a Massive Sponge Die-Off in the Western Mediterranean. Front Microbiol 7:752

10. Simister R, Taylor MW, Tsai P, et al (2012) Thermal stress responses in the bacterial biosphere of the Great Barrier Reef sponge, *Rhopaloeides odorabile*. Environ Microbiol 14:3232–3246

11. Pita L, Erwin PM, Turon X, López-Legentil S (2013) Till death do us part: stable sponge-bacteria associations under thermal and food shortage stresses. PLoS One 8:e80307

12. Strand R, Whalan S, Webster NS, et al (2017) The response of a boreal deep-sea sponge holobiont to acute thermal stress. Sci Rep 7:1660

13. Li CW, Chen JY, Hua TE (1998) Precambrian sponges with cellular structures. Science 279:879–882

14. Love GD, Grosjean E, Stalvies C, et al (2009) Fossil steroids record the appearance of Demospongiae during the Cryogenian period. Nature 457:718–721

15. Bell JJ (2008) The functional roles of marine sponges. Estuar Coast Shelf Sci 79:341–353

16. Bergquist PR (2001) Porifera (Sponges). In: eLS. John Wiley & Sons, Ltd

17. Hooper JNA, Van Soest RWM (2002) Systema Porifera. A Guide to the Classification of Sponges. In: Hooper JNA, Van Soest RWM, Willenz P (eds) Systema Porifera. Springer US, pp 1–7

18. Taylor MW, Hill RT, Piel J, et al (2007) Soaking it up: the complex lives of marine sponges and their microbial associates. ISME J 1:187–190

19. Southwell MW, Weisz JB, Martens CS, Lindquist N (2008) *In situ* fluxes of dissolved inorganic nitrogen from the sponge community on Conch Reef, Key Largo, Florida. Limnol Oceanogr 53:986

20. Fiore CL, Baker DM, Lesser MP (2013) Nitrogen biogeochemistry in the Caribbean sponge, *Xestospongia muta* : a source or sink of dissolved inorganic nitrogen? PLoS One 8:e72961

21. Muscatine L, Porter JW (1977) Reef Corals: Mutualistic Symbioses Adapted to Nutrient-Poor Environments. Bioscience 27:454–460

22. Taylor MW, Radax R, Steger D, Wagner M (2007) Sponge-associated microorganisms: evolution, ecology, and biotechnological potential. Microbiol Mol Biol Rev 71:295–347

23. Hentschel U, Piel J, Degnan SM, Taylor MW (2012) Genomic insights into the marine sponge microbiome. Nat Rev Microbiol 10:641–654

24. Webster NS, Botté ES, Soo RM, Whalan S (2011) The larval sponge holobiont exhibits high thermal tolerance. Environ Microbiol Rep 3:756–762

25. Erwin PM, Pita L, López-Legentil S, Turon X (2012) Stability of sponge-associated bacteria over large seasonal shifts in temperature and irradiance. Appl Environ Microbiol 78:7358–7368

26. Morrow KM, Bourne DG, Humphrey C, et al (2015) Natural volcanic CO2 seeps reveal future trajectories for host-microbial associations in corals and sponges. ISME J 9:894–908

27. Galitz A, de C. Cook S, Ekins M, et al (2018) Identification of an aquaculture poriferan “Pest with Potential” and its phylogenetic implications. PeerJ 6:e5586

28. Pichler M, Coskun ÖK, Ortega-Arbulú A-S, et al (2018) A 16S rRNA gene sequencing and analysis protocol for the Illumina MiniSeq platform. Microbiology Open 215:e00611

29. Rognes T, Flouri T, Nichols B, et al (2016) VSEARCH: a versatile open source tool for metagenomics. PeerJ 4:e2584

30. Oksanen J, Blanchet FG, Friendly M, et al (2017) vegan: Community Ecology Package

31. Love MI, Huber W, Anders S (2014) Moderated estimation of fold change and dispersion for RNA-seq data with DESeq2. Genome Biol 15:550

32. Wilson JB (1991) Methods for fitting dominance/diversity curves. J Veg Sci 2:35–46

33. Thomas T, Moitinho-Silva L, Lurgi M, et al (2016) Diversity, structure and convergent evolution of the global sponge microbiome. Nat Commun 7:11870

34. Frontier S (1987) Applications of Fractal Theory to Ecology. In: Developments in Numerical Ecology. Springer Berlin Heidelberg, pp 335–378

35. Wilson JB, Terry C. E. Wells, Trueman IC, et al (1996) Are there Assembly Rules for Plant Species Abundance? An Investigation in Relation to Soil Resources and Successional Trends. J Ecol 84:527–538

36. Bell JJ, Davy SK, Jones T, et al (2013) Could some coral reefs become sponge reefs as our climate changes? Glob Chang Biol 19:2613–2624

37. Bell JJ, Bennett HM, Rovellini A, Webster NS (2018) Sponges to be winners under near-future climate scenarios. Bioscience 68:955–968

