## Supplementary Tables for "Short-term exposure to high-water temperature causes a shift in the microbiome of the common aquarium sponge *Lendenfeldia chondrodes*"

**Supplementary Table 1:** *Lendenfeldia chondrodes* core OTUs and their associated taxonomic affiliation according to the Silva 16S rRNA database.

| OTU name | Silva Classification |
|---|---|
| CBAS_OTU_1 | Bacteria;Cyanobacteria;Oxyphotobacteria;Synechococcales;Cyanobiaceae;[Synechococcus] spongiarum group |
| CBAS_OTU_2 | Bacteria;Bacteroidetes;Rhodothermia;Rhodothermales;Rhodothermaceae;uncultured |
| CBAS_OTU_3 | Bacteria;Proteobacteria;Alphaproteobacteria;Rhodobacterales;Rhodobacteraceae;Ruegeria |
| CBAS_OTU_4 | Bacteria;Proteobacteria;Gammaproteobacteria;Oceanospirillales;Pseudohongiellaceae;Pseudohongiella |
| CBAS_OTU_5 | Bacteria;Cyanobacteria;Oxyphotobacteria;Chloroplast |
| CBAS_OTU_6 | Bacteria;Actinobacteria;Acidimicrobiia;Microtrichales;Microtrichaceae; Sva0996 marine group |
| CBAS_OTU_8 | Bacteria;Proteobacteria;Gammaproteobacteria;Pseudomonadales;Pseudomonadaceae;Pseudomonas |
| CBAS_OTU_11 | Bacteria;Acidobacteria;Subgroup 9 |
| CBAS_OTU_13 | Bacteria;Planctomycetes;Planctomycetacia;Pirellulales;Pirellulaceae;Pir4 lineage |
| CBAS_OTU_18 | Bacteria;Proteobacteria;Gammaproteobacteria;Steroidobacterales;Woeseiaceae;JTB255 marine benthic group |
| CBAS_OTU_21 | Bacteria;Proteobacteria;Gammaproteobacteria;UBA10353 marine group |
| CBAS_OTU_47 | Bacteria;Proteobacteria;Gammaproteobacteria;UBA10353 marine group |
| CBAS_OTU_48 | Bacteria;Proteobacteria;Gammaproteobacteria;Betaproteobacteriales; Burkholderiaceae;Ralstonia |
| CBAS_OTU_50 | Bacteria;Proteobacteria;Gammaproteobacteria;KI89A clade |
| CBAS_OTU_59 | Bacteria;Proteobacteria;Alphaproteobacteria;Rhodovibrionales;Kiloniellaceae;uncultured |
| CBAS_OTU_89 | Bacteria;Bacteroidetes;Bacteroidia;Cytophagales;Cyclobacteriaceae |
| CBAS_OTU_388 | Bacteria;Proteobacteria;Alphaproteobacteria;Rhodobacterales;Rhodobacteraceae |

**Supplementary Table 2:** Mean deviance, standar error deviance and mean Akaike Information Criterion for five Rank-Abundance models applied to Phylum-level data of the *Lendenfeldia chondrodes* microbiome. The analysis of model deviance is presented below together with a post-hoc (Tukey) comparison of the different model fitted to the data.

| | Null | Preemption | Lognormal | Zipf | Mandelbrot |
|---|---|---|---|---|---|
| Meand deviance | 9114.416 | 765.0778 | 1818.8832 | 2620.4862 | NA |
| SE deviance | 1106.230 | 142.3804 | 362.4572 | 490.3839 | NA |
| Mean AIC | 9185.999 | 838.6612 | 1894.4666 | 2696.0696 | NA |

| | Df | Sum Sq | Mean Sq | F value | Pr(>F) |
|---|---|---|---|---|---|
| model | 4 | 95.65 | 23.912 | 34.79 | <2e-16 |
| Residuals | 94 | 64.61 | 0.687 | | |

| | **Preemption** | Lognormal | Mandelbrot | Zipf | Null |
|---|---|---|---|---|---|
| Preemption | | 0.79 | 0.17 | 1.22 | 2.56 |
| Lognormal | 0.027 | | 0.96 | 0.43 | 1.77 |
| **Mandelbrot** | **0.965** | 0.004 | | 1.40 | 2.74 |
| Zipf | 0.0001 | 0.469 | < 0.0001 | | 1.34 |
| Null | < 0.0001 | < 0.0001 | < 0.0001 | < 0.0001 | |

**Supplementary Table 3:** Mean deviance, standard error deviance and mean Akaike Information Criterion for five Rank-Abundance models applied to OTU-level data of the *Lendenfeldia chondrodes* microbiome. The analysis of model deviance is presented below together with a post-hoc (Tukey) comparison of the different models fitted to the data.

| | Null | Preemption | Lognormal | Zipf | Mandelbrot |
|---|---|---|---|---|---|
| Meand deviance | 32467.790 | 16453.30 | 2114.3692 | 1190.8433 | 968.6425 |
| SE deviance | 4306.957 | 2302.41 | 286.3831 | 247.8633 | 144.6762 |
| Mean AIC | 33184.136 | 17171.64 | 2834.7147 | 1911.1888 | 1690.9880 |

| | Df | Sum Sq | Mean Sq | F value | Pr(>F) |
|---|---|---|---|---|---|
| model | 4 | 206.74 | 51.68 | 99.37 | <2e-16 |
| Residuals | 95 | 49.41 | 0.52 | | |

| | Preemption | Lognormal | **Mandelbrot** | Zipf | Null |
|---|---|---|---|---|---|
| Preemption | | 2.00 | 2.81 | 2.68 | 0.69 |
| Lognormal | < 0.0001 | | 0.81 | 0.69 | 2.69 |
| Mandelbrot | < 0.0001 | 0.0051 | | 0.13 | 3.51 |
| **Zipf** | < 0.0001 | 0.0271 | **0.9809** | | 3.38 |
| Null | 0.0025 | < 0.0001 | < 0.0001 | < 0.0001 | |

**Supplementary Table 4:** Permutation test (999 permutations) for a canonical correspondence analysis of the effect of temperature treatment, sponge color morph and their interaction on the composition of the microbiome of *Lendenfeldia chondrodes*. Terms added sequentially, first to last. For this analysis, counts were corrected using the bacterial load of each sample and OTUs with less than 50 counts were removed from the analysis.

| | Df | Chi-Square | F | Pr(>F) |
|---|---|---|---|---|
| Treatment | 1 | 0.36522 | 7.9291 | 0.001 |
| Color morph | 1 | 0.20193 | 4.3840 | 0.002 |
| Treatment x Color morph | 1 | 0.09383 | 2.0370 | 0.071 |
| Residual | 12 | 0.55273 | | |
