## Supplementary Figures for "Short-term exposure to high-water temperature causes a shift in the microbiome of the common aquarium sponge *Lendenfeldia chondrodes*"

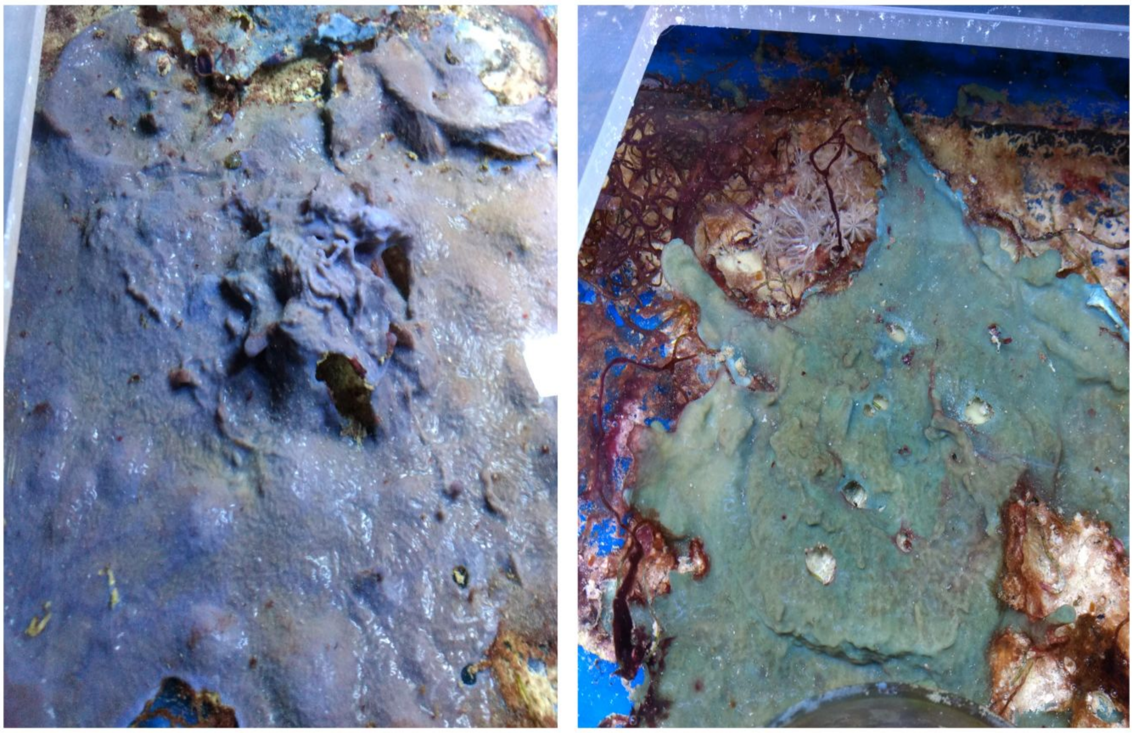

**Supplementary Figure 1:** Purple (left) and green morphs of *Lendenfeldia chondrodes*. Picture not at scale. As a reference, the specimen on the left is ca. 20 cm and the specimen on the right is ca. 10 cm across.

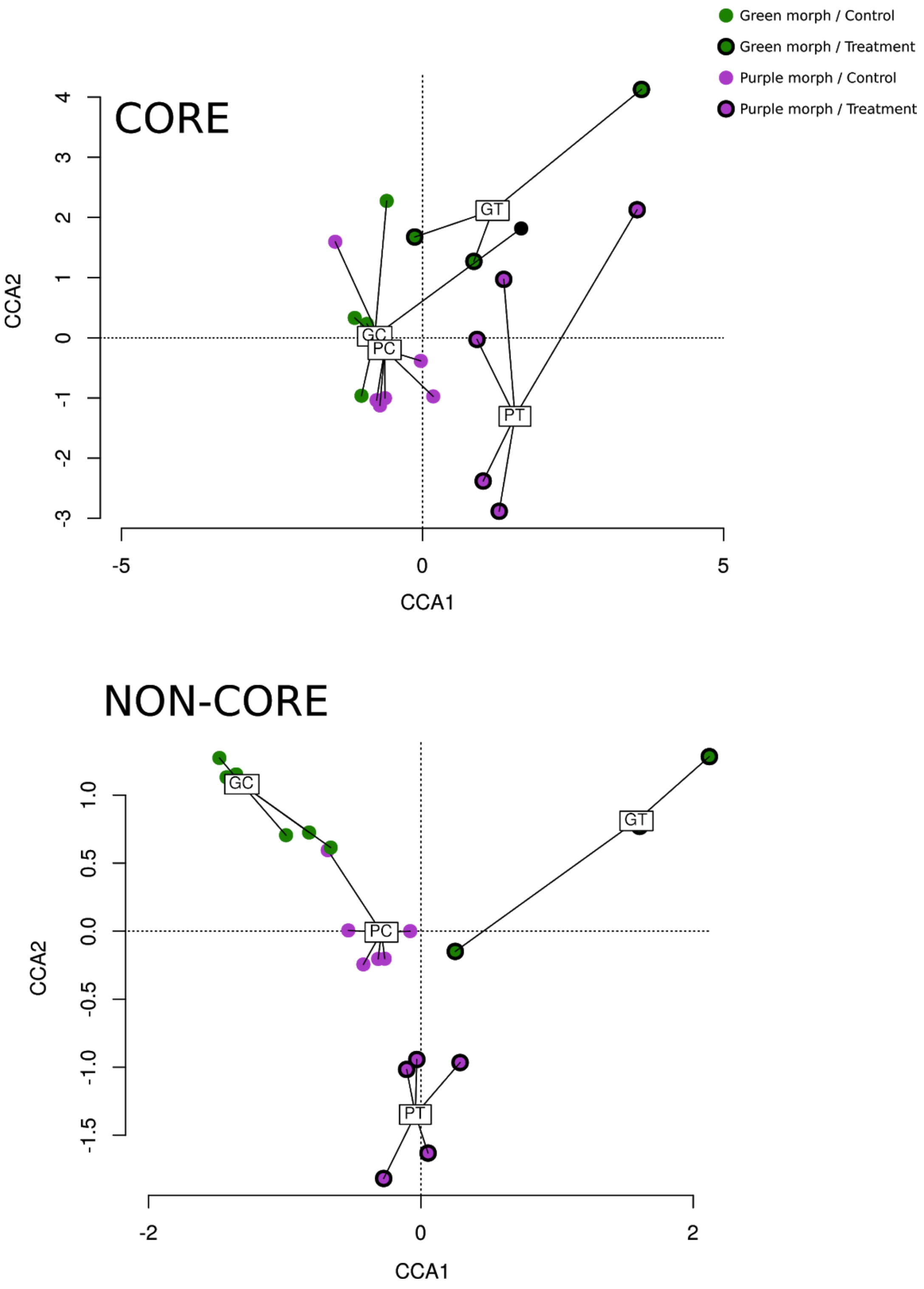


**Supplementary Figure 2:** Canonical correspondence analysis of OTUs belonging to the core and non-core community of *Lendenfeldia chondrodes*.

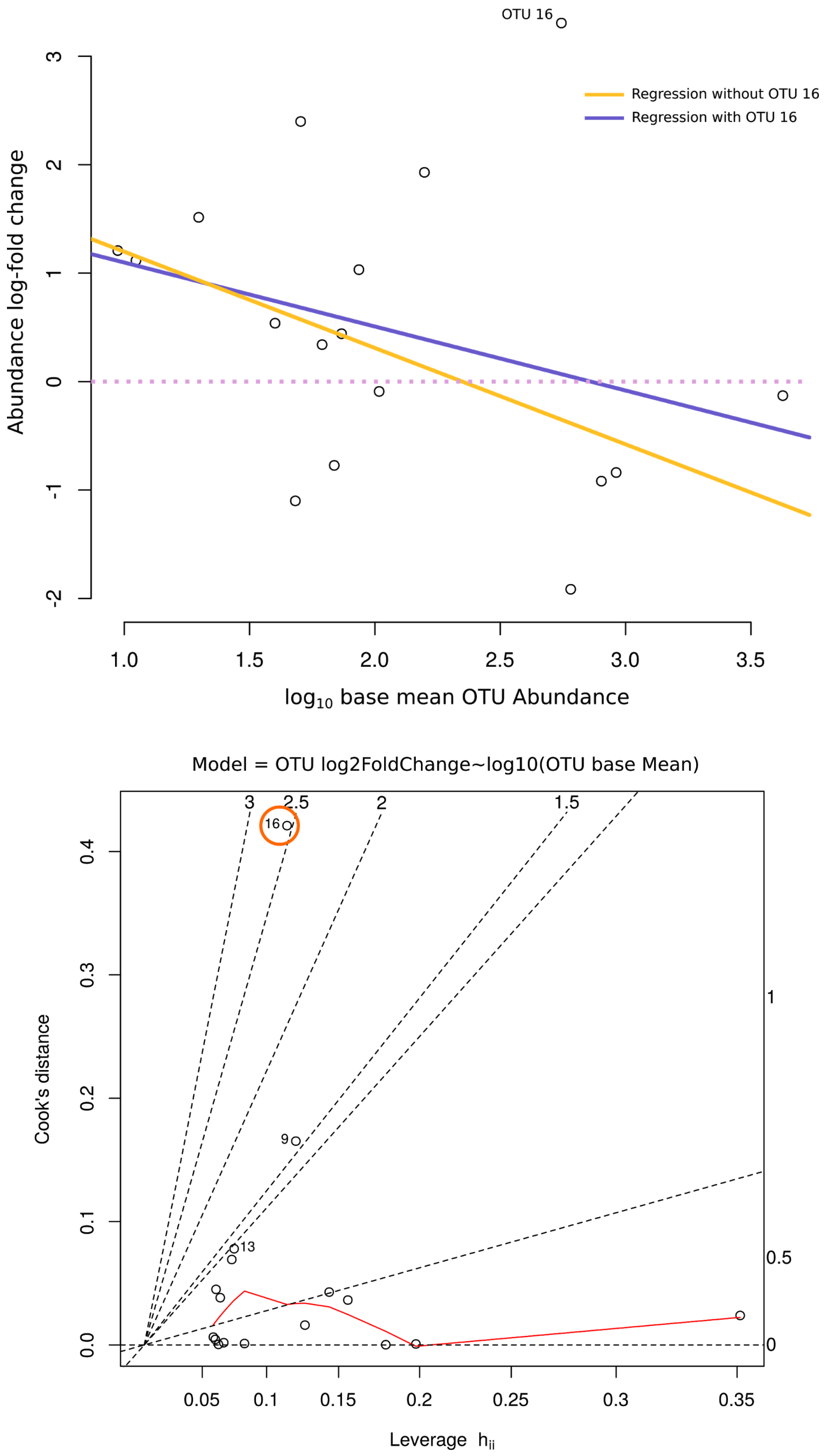


**Supplementary Figure 3:** (Upper panel) Regression analysis of OTU abundance log2-fold change using as predictor the base mean OTU abundance (log10 transformed). The regression line including OTU 16 is shown in purple and OTU 16 is labeled (on the top). The regression excluding OTU 16 is shown in yellow. (Lower panel) Cook's D vs. leverage analysis showing OTU 16 as a point with high influence on the regression's slope.
